# Multiparameter stimulation mapping of signaling states in single pediatric immune cells reveals heightened tonic activation during puberty

**DOI:** 10.1101/2022.11.14.516371

**Authors:** Rohit Farmer, Richard Apps, Juan Quiel, Brian A. Sellers, Foo Cheung, Jinguo Chen, Amrita Mukherjee, Peter J. McGuire, John S. Tsang

**Affiliations:** Trans-NIH Center for Human Immunology, Autoimmunity, and Inflammation, NIH, Bethesda, Maryland, USA; Metabolism Infection and Immunity Section, National Human Genome Research Institute, NIH, Bethesda, Maryland, USA; Multiscale Systems Biology Section, Laboratory of Immune System Biology, National Institute of Allergy and Infectious Diseases, NIH, Bethesda, Maryland, USA

## Abstract

Cellular stimulation via factors such as cytokines followed by multiparameter single-cell measurements is a powerful approach to interrogate cellular functions. However, transforming such high-dimensional data into biological insights presents unique challenges, particularly given the extensive response heterogeneity among single cells, such as the presence of bimodal responding versus non-responding subpopulations upon stimulation. Here we present an unsupervised **h**igh-**d**imensional approach for analyzing **stim**ulation responses at the single cell level (**HDStIM)** and apply it to evaluate how pediatric development may shape peripheral immune cell signaling states and responsiveness to stimulations in 42 subjects (age: 2 - 16). We show that in comparison to the conventional approach of assessing one marker at a time by averaging across single cells, HDStIM can effectively learn, in an unsupervised fashion, the multi-parameter signature of responding versus non-responding cells to accurately quantify responses within cell populations. HDStIM reveals that the extent of pre-stimulation/baseline activation of interferon-related and TCR signaling molecules in myeloid and T cells, respectively, increases during puberty. This suggests that puberty is marked by a heightened “tonic” activation state in these cells, perhaps to strengthen defense against pathogens during this period of human development.

## Introduction

Experiments with a cellular stimulus combined with monitoring of a large number of response variables at the single cell level are becoming increasingly prevalent (Aghaeepour et al., 2017; Arunachalam et al., 2020; Bendall et al., 2011). A defining feature of the resulting data is that the extensive heterogeneity of responses observed at the single cell level (Buettner et al., 2015; Efremova et al., 2020; Satija and Shalek, 2014), revealing a continuity of phenotypes that can blur the distinctions between cell types and cell states (Morris, 2019). These observations challenge the notion that cellular responses and their states can be analyzed and quantified one marker at a time by averaging across single cells, for example, as is often done in the analysis of the activation status of intracellular signaling molecules after cytokine stimulation of immune cells (Bendall et al., 2011). Such single marker, cell-averaging approaches can mask important information about response heterogeneity and cell states (e.g., stimulation often induces responses from only a subset of cells). Janes et al. have assessed multiple variables for contribution to responses in the context of tumour necrosis factor stimulation using regression models, but this is limited by a priori definition of a response phenotype (Janes et al., 2005). Here we present an approach (called **HDStIM**; implemented in an R package) that utilizes multiple parameters (e.g., phosphorylation status) simultaneously to assess responses at the single cell level in an unsupervised manner. HDStIM can utilize all markers observed or a subset of them selected a priori to identify the cell subpopulations responding specifically to the stimulation. HDStIM infers the response phenotype defined by multiple markers and provides a quantitative sense of the relative contribution of these markers to defining the response phenotype.

We designed and tested HDStIM using data from stimulation assays where mass cytometry was used to measure signaling responses by immune cells after brief (∼15 minutes) in vitro stimulations with cytokines and other immune ligands (Fernandez and Maecker, 2015). Such cytometry panels typically comprised ∼20 surface protein markers for identifying broad immune cell lineages and subpopulations, as well as ∼10-20 markers of intracellular signaling responses that predominantly mark phosphorylation states of intracellular signaling proteins. We expect HDStIM to be applicable to related assays, e.g., measurements by high parameter flow cytometry (Saeys et al., 2016) and single cell multiomics such as CITE-seq (Stoeckius et al., 2017). We first applied HDStIM to a previously published dataset to demonstrate that HDStIM could improve discrimination of responding cells, delineating the characteristics of scenarios under which our high dimensional analysis can identify responses and their signatures not detected by conventional assessment of individual markers by averaging across single cells (Bendall et al., 2011). Taking advantage of HDStIM’s capabilities, we next applied it to a new dataset we generated from a pediatric population (N=42; ages 2 - 16) to assess how peripheral immune cells and their responses to stimulations may be modulated during development. While peripheral immune cell phenotypes have been assessed as a function of age, the intracellular signaling statuses before and after stimulation have not been well characterized throughout pediatric development. HDStIM reveals associations with age and specifically highlights puberty as an immunologically distinct period of development. Specifically, the extent of pre-stimulation/baseline activation of intracellular interferon-related and TCR signaling molecules in myeloid and T cells, respectively, increases substantially before trending down after puberty. These observations suggest that puberty is marked by a heightened “tonic” activation state in the immune system, perhaps to strengthen defense against pathogens during this period of human development.

## Results

### Unsupervised learning of response phenotypes and subpopulations

HDStIM utilizes data from both unstimulated and stimulated samples for which multiple response markers are assayed. The in vitro stimulation datasets we analyzed were generated in multi-well plates covering multiple experimental stimulation conditions involving a mix of cell types - in these instances, human peripheral blood mononuclear cells (PBMCs). Stimulations were short (∼15 minutes) and standardized across all wells designed to assess early signaling responses without interference from feedforward regulation or overt interference by secreted factors from other cells. Mass cytometry panels approaching 40 markers can include both protein markers for population identification as well as protein-modification markers of signaling states such as pathways responding to TCR, LPS, or cytokine stimulation. Specific cell populations and subsets were subsequently identified by manual gating or automated clustering of cells pooled from pre-stimulation (baseline) and stimulation conditions. Such a dataset can be viewed as a matrix comprising cell populations intersecting with (stimulated and unstimulated) conditions across different stimuli. Response phenotypes, e.g., the phosphorylation status of the signaling proteins measured, can then be linked to each combination of cell population and stimulation condition (“stimulation-population combinations,” or SPC).

To develop HDStIM, we hypothesize that any stimulation would result in a subpopulation of “responding” cells with a distinct response phenotype defined by multiple markers (e.g., phosphorylation of signaling molecules). This responding subpopulation can be as large as the entire stimulated cell population, but as existing data suggest, it can also be a subset since not all cells respond or some cells only respond weakly to certain stimuli. Our goal is to take advantage of this heterogeneity and simultaneously utilize the unstimulated/baseline and stimulated samples to learn, in an unsupervised manner in high-dimensional space, what the response phenotype is for a given SPC **(Figure 1A)**. Thus, for each SPC by using both experimentally stimulated and unstimulated cells, HDStIM first performs unsupervised clustering using all response markers to partition single cells into two groups comprising the responding and non-responding subpopulations. Our second hypothesis is that cells with the response phenotype are more prevalent in frequency and thus are enriched in the stimulated condition. While some cells at baseline (before stimulation) may already have a phenotype that resembles the responding population, our assumption is that the stimulation is sufficient to induce a higher frequency of cells with that phenotype. Thus, for each of the two cell clusters identified above, HDStIM tests to assess which one of the two clusters has a significantly higher frequency in the experimentally stimulated condition (by using Fisher’s exact test with a nominal cutoff: p<0.05) and labels this as the “responding” population. An insignificant Fisher’s exact test would suggest that the experimental stimulation is unable to generate a sufficiently distinct response phenotype (e.g., due to a weak stimulus or noise in the experiments), otherwise we consider HDStIM able to identify the responding population. A feature of this approach is that it naturally identifies “responding” cells in the unstimulated (baseline) condition as well: these are cells that have the responding phenotype prior to stimulation. HDStIM quantifies the response to the stimulation as the increase in the frequency of the responding cell population in the stimulated vs. the unstimulated (baseline) conditions **(Figure 1A)**.

**Figure 1:**
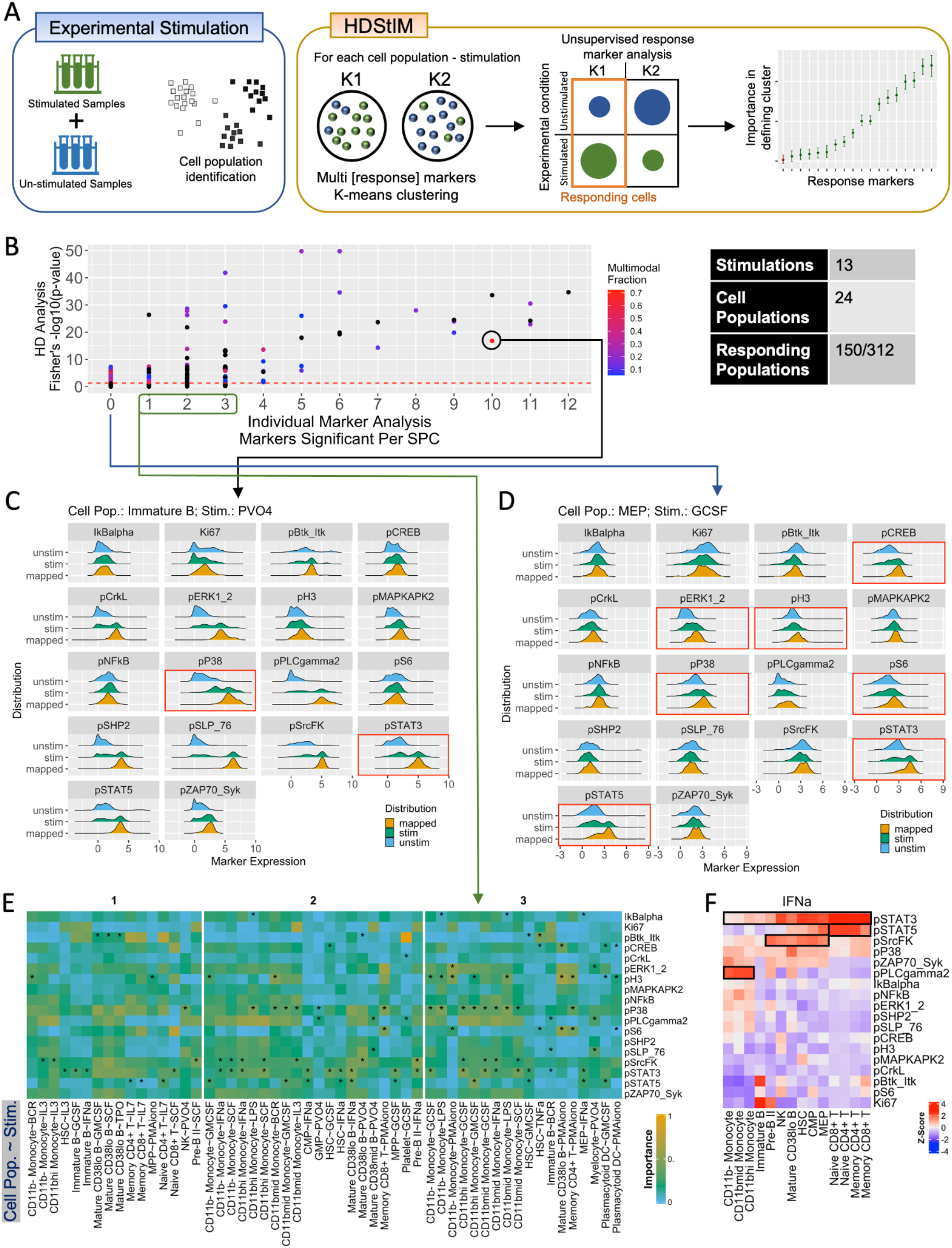
Validation of HDStIM using a previously published bone marrow cell dataset. A) Schematic outline for the use of multiple response markers by HDStIM to identify cells with a responding phenotype from unstimulated and stimulated experimental samples for a given SPC. B) HDStIM was compared to the conventional marker by marker analysis from a previously published dataset. For all 312 SPCs the number of response markers significant in individual marker analysis (x-axis), is plotted against the probability for identification of a responding population by HDStIM, with color representing the fraction of response markers with multimodal distribution. C) HDStIM improved the identification of responses by a fraction of a cell population. In this example for immature B cells stimulated by pervanadate (PV04), all 18 response markers are shown for experimentally unstimulated (blue), stimulated samples (green), or the responding population identified by HDStIM (orange). Markers p38 and STAT3, boxed red, highlight the distinct multimodal response detected by HDStIM. D) HDStIM was able to identify a responding fraction where modest responses occur in multiple markers. This example for megakaryocyte-erythroid progenitor cells stimulated by GCSF, with plots as described in panel C, shows markers responsible (boxed red). E) HDStIM identified additional response markers of importance for the definition of a responding population. 54 SPCs are shown, with colors indicating the importance of each response marker for the definition of the responding population, where in each case HDStIM identified at least one response marker ranked higher than those identified by asterisks which were significant in individual marker analysis. (F) For all SPCs with experimental stimulation by IFNa in which HDStIM identified a responding population, the relative ranking for contribution to this responding fraction is plotted for all response markers with populations clustered by their response phenotypes. Responses to IFNa segregate into 3 groups defined by increasing responses to STAT3 combined with responses by either PLCg2, SrcFK, or STAT5.

While HDStIM uses all signaling markers to infer the responding phenotype and subpopulation, individual markers contribute by different quantitative extents to define the response phenotype and it is therefore biologically informative to know which markers contributed the most and are the most important markers defining the response phenotype. Towards this end, for each SPC a Random Forest (RF) based machine learning approach is used by HDStIM to classify cells between the responding and non-responding cell clusters with the goal to learn from the RF classifier the “importance score” of each responding marker (see Methods; **Figure 1A)**. This allows HDStIM to include all markers for the determination of the responding phenotype while reporting the markers that contributed the most for defining the responding population for each SPC.

### Assessing HDStIM against single marker analysis

We first applied HDStIM to a previously published dataset in which bone marrow cells from a single human donor were stimulated *in vitro* with 13 stimuli and the responses were monitored using 18 intracellular markers in 24 cell populations (Bendall et al., 2011) **(Supplementary Tables 1-3)**. For the 312 SPCs (13 stimuli x 24 cell populations), the Fisher’s exact test p values (i.e., confidence for the identification of a responding population by HDStIM) are shown in **Figure 1B**: 150 SPCs are nominally significant at p<0.05 and these are contrasted with the result from the previously published single marker analysis based on the average fold-change across single cells in the stimulated vs. the unstimulated conditions. This comparison shows that HDStIM was able to identify a responding population for all SPCs in which individual maker-based tests found at least four markers to be significant. This is consistent with our expectation that HDStIM is the most effective when the responding population is characterized by more than one marker. HDStIM also detected a responding population for 66% of the SPCs (69/104) where 1-3 response markers were found significant by the individual marker-based tests. Importantly, HDStIM was able to identify a responding population in 54 out of 181 SPCs where individual marker analysis did not detect a single significant marker.

We next sought to better understand the scenarios under which HDStIM can improve the detection of responding populations in comparison to individual marker analysis. Notably as mentioned above, HDStIM can achieve better sensitivity by identifying a subset of cells that respond in a given SPC; such bimodality in responses is a prevalent feature in this dataset and supports our hypothesis that upon stimulation typically only a subset of cells would show a detectable response phenotype. Of the 150 SPCs for which a responding population was found, 94 have multimodal distribution for at least one response marker, and this was robust to the P-value threshold used for establishing deviation from a unimodal distribution **(Figure 1B and Supplementary Figure 1A)**. That HDStIM can effectively identify a subset of responding cells is illustrated by the example of pervanadate stimulated immature B cells, where responses in p38 and STAT3 in particular were multimodal **(Figure 1C)**. Comparing the fold-changes in median intensity between stimulated and unstimulated cells for all immature B cells would underestimate the response for both of these markers, whereas HDStIM was able to pinpoint the subset of immature B cells that responded to pervanadate to enable a more accurate definition and quantitation of the response.

A second scenario under which HDStIM differs significantly from single marker analysis is when modest responses occur in multiple markers such that HDStIM can identify a responding cell fraction even though no individual markers show a sufficiently significant change. In total, HDStIM identified responding populations for 54 SPCs where no individual markers showed a significant response. An example of this involved the GCSF-stimulated megakaryocyte-erythroid progenitor cells: HDStIM revealed that the responding cells were marked by the coordinated responses of at least seven markers **(Figure 1D)**. HDStIM could thus potentially help reveal cell-type specific crosstalk among signaling pathways that involve multiple signaling markers.

A related, third scenario under which HDStIM differed from single marker analysis was by implicating additional response markers when defining the responding cell subset. We observed 54 SPCs in which individual maker analysis showed 1–3 significant response markers, while at least one additional marker was ranked higher based on the variable importance score provided by Random Forest, suggesting that these additional markers might be biologically relevant as they have contributed to the identification of the responding fraction **(Figure 1E)**. Overall, for all 150 SPCs where HDStIM detected a responding population, the variable importance scores, as expected, showed significant negative correlation with the adjusted p-values from individual marker analysis (r = -0.22, p-value = 0.003). However, the modest strength of this correlation is consistent with the detection of potential new markers of the responding population/phenotype by HDStIM that were not detected by single marker analysis **(Supplementary Figure 1B)**.

The importance scores for individual markers provide biological insights of the simulation response. For example, signaling responses to IFN stimulation appeared cell type specific. IFNa stimulated signaling response profiles clustered into three major cell groups comprising monocytes, immature/progenitor cells (including hematopoietic stem cells), and T cells. These three cell clusters were defined by common responses in STAT3 in combination with, respectively, PLCg2, SrcFK, or STAT5 **(Figure 1F)**. To the best of our knowledge, PLCg2 in monocytes and SrcFK in progenitor cells are potentially novel cell-type specific signaling mediators of IFNa responses (Ivashkiv and Donlin, 2014; Jefferies, 2019). Notably, these were only detected when using HDStIM but not when markers were analyzed individually or when experimentally stimulated data were assessed alone without pooling data from the baseline/pre-stimulation condition as is performed by HDStIM by default **(Supplementary Figure 1C & 1D)**. Together these observations indicate that HDStIM can leverage information from multiple markers to identify responding cells, pinpoint the responsible markers, and provide additional biological insights in comparison to conventional single marker analysis.

### Analysis of stimulation responses at single-cell resolution in pediatric subjects spanning puberty

We next sought to apply HDStIM to assess how the signaling response to stimuli in human immune cells change as a function of pediatric development. Increasing evidence suggests profound shifts in immune system functions and responsiveness during early to adolescent development due to genetic, developmental, and cumulative exposure factors (Kane and Ismail, 2017; Klein and Flanagan, 2016; Lamason et al., 2006). However, the signaling response in primary immune cells across the pediatric age range have not been well studied using unbiased systems biology approaches. Here we assessed the stimulation responses of single PBMCs from a pediatric cohort of 42 subjects ranging in age from 2 to 16 years old with even distribution between the two sexes **(Figure 2A)**. Each PBMC sample was stimulated with 4 stimuli (IFNa, TCR, LPS, IFNg) and the responses were monitored for 10 intracellular signaling markers in 19 cell populations identified from unbiased clustering of the single cell mass cytometry data obtained from before and after stimulation (see Methods and **Supplementary Tables 1-3**). Of the 76 SPCs (4 stimuli x 19 cell populations) HDStIM identified 28 SPCs with responding subpopulations. The markers that contributed significantly to defining these responding populations are largely consistent with existing knowledge about the underlying signaling pathways, for example, IFNa and IFNg induced STAT responses, while TCR stimulation activated ERK and AKT **(Figure 2B)**.

**Figure 2:**
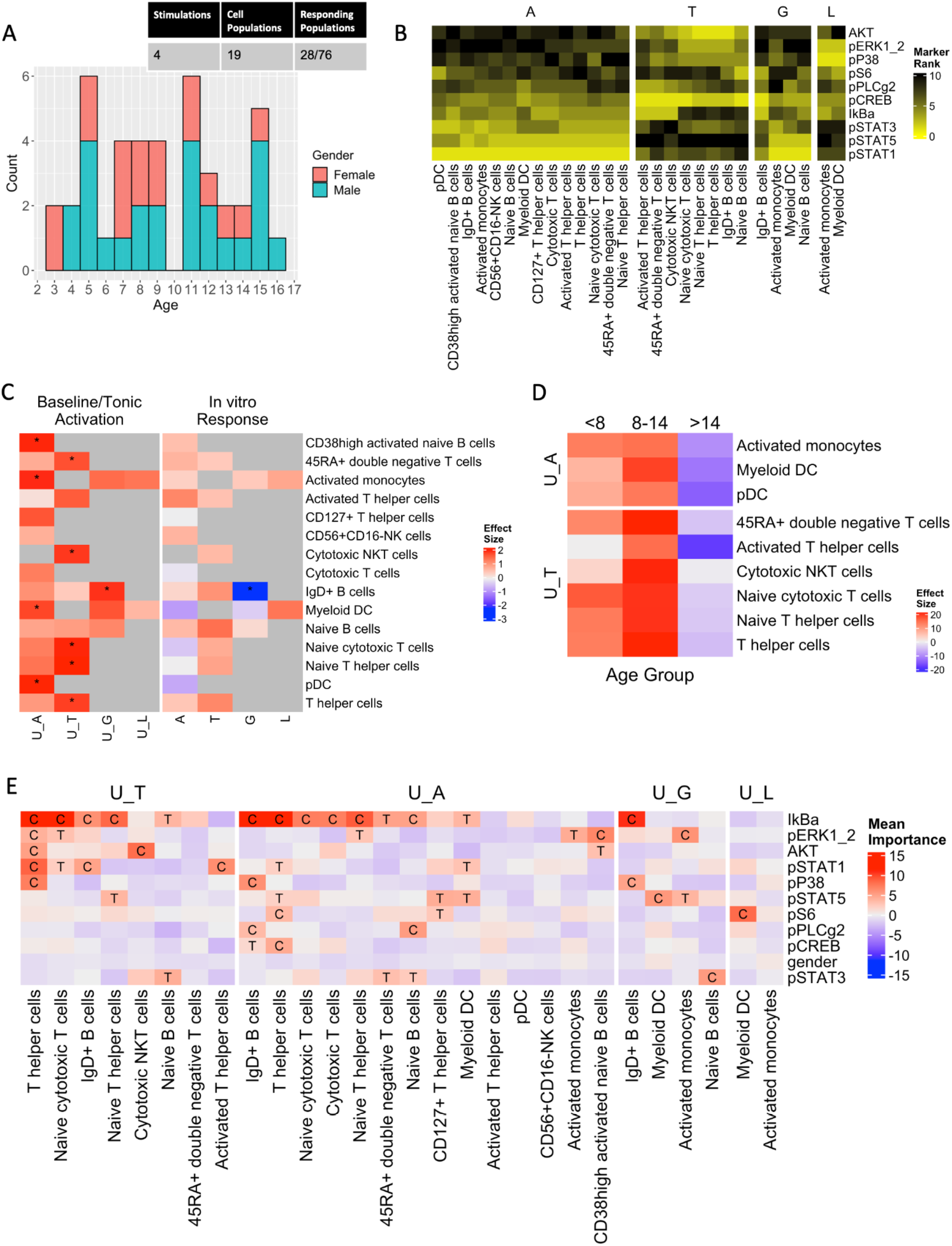
Application of HDStIM using a novel PBMC dataset reveals effects of pediatric age on immune responsiveness. A) Distributions of age and sex in this dataset support assessment of their effect on stimulation responses throughout childhood. B) Analysis of in vitro responses using HDStIM identified responding populations for 28 SPCs, for which mean importance scores are shown for all response markers grouped by stimulus where A indicates IFNa, G indicates IFNg, T indicates TCR and L indicates LPS. C) For all 28 SPCs the effect of age on either the fractions of cells with baseline/tonic activation (left), or the increase in this fraction in response to in vitro stimulation (right), was assessed with plots showing estimates and P-values for the effect of age from linear modeling. SPCs with P-values below 0.05 are marked with an asterisks. The experimental stimulations used to define each baseline/tonic activation phenotype assessed are labeled as “U_[stimulant]”. D) Increases in baseline activation that were observed with age throughout childhood, were further analyzed for the subgroups of age <8, 8-14, or >14. For fractions of cells identified as responding at baseline based on IFN-a stimulation for all myeloid cells (top) and based on TCR stimulation for all T-cell populations (bottom), size estimates are plotted for the effect of age in spline regression modeling. E) To address the molecular basis of the changes in stimulation responses with age, for all 28 stimulation-cell populations where an responding population was identified, the fraction of cells detected to be responding at baseline was used to assess the magnitude of individual response markers for correlation with age. Relative importance scores are plotted for each marker with ‘C’ indicating confirmed, or ‘T’ indicating a tentative effect; unlabelled markers were deemed unimportant by Boruta (see Methods).

To assess the effects of age and sex on the stimulation responses we considered two parameters that can be extracted for each subject after the identification of responding cells by HDStIM. First, the fraction of cells found to have a responding phenotype in the unstimulated (baseline) experimental condition was used as the level of ‘baseline/tonic activation’, as defined for a given SPC after in vitro stimulation. Second, the difference between this baseline and the fraction of cells found to have a responding phenotype in the stimulated condition was used as a measure of the magnitude of the actual in vitro response for each SPC. For the 28 SPCs where a responding population was identified, these per-subject baseline and response fractions were assessed for the effects of age, gender, or their interaction using linear modeling. Age was the only variable that showed significant associations, particularly involving the extent of baseline tonic activation of both myeloid and T cells (nominal p<0.05) **(Figure 2C)**. In particular, the responding phenotype observed after IFNa stimulation was also found at baseline (in the pre-stimulation condition) to positively associate with age in plasmacytoid dendritic cells (pDCs), conventional/myeloid DCs, activated monocytes (HLA-DR+), and CD38hi naive B-cells. Similarly, the responding phenotype observed after TCR stimulation was also seen at baseline to increase with age in several T-cell populations, including naive CD4+ and CD8+ cells. The response magnitude as defined above, however, did not show any clear associations with age.

Puberty is a period of profound developmental change (Patton and Viner, 2007). We thus wondered if the age associations observed were partly driven by changes linked to puberty. We divided subjects into three groups: before (age <8), during (age 8-14), or after puberty (age >14), and assessed the association between these groups and the baseline/tonic activation and response phenotypes as defined above, but here by using spline regression with “knots” at 8 and 14 years of age (see Methods). For both the myeloid and T-cell baseline tonic activation phenotypes, the strongest positive estimates for the effect of age (compared to the age<8 group as a reference) were for the age 8-14 group, followed by a negative estimate (thus a decline) for the age>14 group. These results suggest that the increases in tonic activation of both myeloid and T cells as a function of age can be mainly attributed to increases associated with puberty **(Figure 2D)**. Our Random Forest based ranking of the responding markers suggests that phosphorylated IkBa contributed the most for defining the baseline activation phenotype; this was true particularly for lymphocytes involving TCR or IFNa stimulation, while pSTAT1 and pSTAT5 were more prominent for myeloid/conventional DCs **(Figure 2E)**. These observations are consistent with established molecular phenotypes of activation responses in T and myeloid cells. IkBa phosphorylation together with increases in ERK and AKT phosphorylation (which were also observed for the baseline responding phenotypes) in T cells may mark a “poised”, more responsive state in these cells. Our observations are also consistent with previous reports of association between pediatric age and longer-term stimulation responses (6-24 hours post stimulation) in T cells (Cohen et al., 2021; Rudolph et al., 2018). A potential function of such heightened tonic activation in antigen-presenting and T cells is to increase baseline defense against infectious pathogens, which exert a heavy toll in childhood mortality and can potentially derail normal physiological and sexual development (Black et al., 2003). Thus, by integrating information from unstimulated and stimulated cells from a pediatric cohort, HDStIM identified a cell type dependent, multi-parameter tonic signaling signature that is associated with puberty in humans.

## Discussion

Here we present HDStIM to leverage the high-resolution landscapes of cellular responses detected in stimulation experiments followed by single-cell measurements. We demonstrate that HDStIM can utilize information from multiple markers in higher dimensional space to identify the responding cell fraction and the relative contribution of markers to these response phenotypes. This is partly enabled by simultaneous, unsupervised analysis of single cell data from both unstimulated and stimulated cells. Short *in vitro* stimulation experiments with measurement of responses by high-parameter cytometry are particularly well suited to analysis by HDStIM. In the context of these datasets, we defined several scenarios under which HDStIM identified responses that could not be detected by conventional median-based analysis of markers individually. HDStIM does not perform averaging across single cells but instead leverages the heterogeneity among single cells to identify subsets of responding cells for a given SPC. HDStIM can pinpoint responding cell fractions even when the response is modest at the individual marker level; it can also implicate additional response markers in comparison to individual response marker analysis. For any SPC in which a responding population can be defined by HDStIM, the variable importance scores that HDStIM reports (based on Random Forest learning) for individual markers were demonstrated to provide biological insight regarding the stimulation responses. For example, when re-analyzing IFNa stimulated responses in bone marrow cells, the importance scores identified PLCg2 and SrcFK as potential cell type specific, novel signaling mediators of responses in monocytes and several types of progenitor cells, respectively.

HDStIM was then applied to link signaling states in immune cells to pediatric development. At ages corresponding to puberty this revealed increased pre-stimulation/baseline activation of intracellular interferon-related and TCR signaling molecules in myeloid and T cells, respectively. A heightened “tonic” activation state in the immune system may strengthen defense against pathogens, but could concomitantly associate with harmful consequences of higher grades of inflammation and even allergy or autoimmunity (Kotliarov et al., 2020).

Temporally specific adoption of the heightened state during puberty may help to balance these opposing consequences. Notably, HDStIM revealed that it was the fraction of populations with a responding phenotype at baseline (in unstimulated cells) – with the responding phenotype solely defined/guided by the phenotype observed after in vitro stimulation– rather than the increase in the frequency of responding cells after in vitro stimulation that we have demonstrated to be associated with puberty. This was similar to an earlier observation in pregnant women where the levels of phosphorylation in certain signaling markers in unstimulated cells contributed the most to the prediction of gestational age (Aghaeepour et al., 2017). This suggests shifts in endogenous responsiveness to steady-state levels of stimulation may underlie some changes in immunity which occur over time.

The increases in phosphorylation of IkBa in several lymphocyte populations could reduce the constitutive inhibition of NF-κB in these cells, and thus increase endogenous responsiveness of T-cells during puberty (Christian et al., 2016). Stable shifts in immune cell phenotype in adults have been linked to epigenetic reprogramming. A prominent example is BCG vaccination, where a defined change in chromatin organization in monocytes alters *ex vivo* responsiveness to TLR stimulation measured 3 months after vaccination (Kong et al., 2021). Epigenomic screening has also revealed increased frequencies of monocyte populations defined by reduced AP-1 accessibility, at 30 days after vaccination against both seasonal influenzas and with the mRNA vaccine BNT162b2 against SARS-CoV-2 (Arunachalam et al., 2021; Wimmers et al., 2021). It remains to be seen whether epigenetic changes could similarly underlie the alterations in IkBa phosphorylation to increase the constitutive activation of T-cells during puberty, although extensive epigenetic changes associated with puberty have been observed in genome-wide methylation studies (Thompson et al., 2018).

Together our results demonstrate the utility of HDStIM in extracting novel biological insights from single-cell stimulation data. Beyond the cytometry-based stimulation assays assessed here, we propose that a high dimensional definition of responding populations could be valuable in other experimental designs. For the rapidly increasing contexts where a stimulus or perturbation is combined with monitoring of a large number of response variables at the single cell level HDStIM represents a robust, end-to-end unsupervised package for unbiased identification of responding cells and associated phenotypes.

## Limitations of the study

Although carefully conducted, this study may have some limitations; for example, the pediatric cohort has fewer samples for age >14. Multimodality in responses could represent more heterogeneity than just responding vs. non-responding. The effect of a larger number of noisy markers is not yet apparent and needs further experimentation. We have used nominal P-values without multiple testing corrections meant for exploratory analysis to demonstrate the tool’s value, less so to define puberty-associated immune phenotypes definitively. Nonetheless, we uncovered both well-established phenotypes using unbiased analysis (so reassuring) and intriguing observations replicable across different T cell subsets (e.g., IkB phosphorylation).

## Acknowledgements

This research was supported by the Intramural Research Program of the NIH, the National Institute of Allergy and Infectious Diseases, and other institutes supporting the Trans-NIH Center for Human Immunology, Autoimmunity, and Inflammation. The content of this publication does not necessarily reflect the views or policies of the Department of Health and Human Services, nor does the mention of trade names, commercial products, or organizations imply endorsement by the U.S. Government.

## Methods

### Data sets

We used two mass cytometry stimulation assay datasets in the development of HDStIM. First, data from the stimulation assay on bone marrow mononuclear cells (Bendall et al., 2011) was obtained from Cytobank reports (https://reports.cytobank.org/1/v1). Stimulation panels contain 13 phenotyping and 18 response markers **(Supplementary Table 1)**. For clarity, we have only used data for the 13 simulation conditions **(Supplementary Table 2)** from donor 1, excluding any combination with the inhibitors used in the original study. In addition, we used the 24-cell population definition defined by Bendall et al. 2011 exported from Cytobank. Like the original article, the raw expression values were first divided by 5 and then arcsinh transformed before further usage. To test the accuracy of our transformation, we reproduced the one-sample t-tests for donor 1, as mentioned in Table S3 of the original article’s supplementary material (Bendall et al., 2011). Second, we repurposed data from a novel in-house stimulation assay on a pediatric cohort of 42 subjects ranging from 2 to 16 years old, grouped into 25 males and 17 females. Our stimulation panel included 20 phenotyping markers and 10 response markers **(Supplementary Table 1)**. For each of the 42 subjects, we performed 4 different stimulation experiments along with an unstimulated control **(Supplementary Table 2)** - following the CyTOF protocol described by (Fernandez and Maecker, 2015). Finally, we used the CyTOF workflow (Nowicka, Malgorzata et al., 2019) to identify 19 cell populations via automated clustering. Pre-clustering CyTOF workflow divides raw expression values by 5 and then arcsinh transform and uses FlowSOM (Van Gassen et al., 2015) as the clustering algorithm.

Our pediatric cohort was initially part of a study of mitochondrial disease in which 21 cases with genetic defects in the oxidative phosphorylation pathway were recruited, along with 21 age-matched healthy controls. However, likely due to heterogeneity in the diseased subjects including a wide range of mutated genes with varied phenotypes related to oxidative stress that is associated with mitochondrial dysfunction (Mc Guire et al., 2009), we could not establish coherent differences in immune phenotypes between the disease and healthy groups **(Supplementary Figure 2B)**. Therefore, we pooled the data for all 42 subjects together and repurposed it to assess an association between immune responses and age-spanning puberty **(Figure 2C)**. To support that the effects of age observed in Figure 2C were not driven by pooling these sample groups, we split the data into cases and controls and repeated the analysis as in Figure 2C, showing the same trends. For example, baseline activation increased with age for myeloid populations in controls and T cell populations in cases, although fewer effects were significant with the smaller sample numbers **(Supplementary Figure 2A)**. To further confirm that cases and controls did not differ in this dataset, using raw response marker values in contrast to activation statuses determined by HDStIM, we utilized unbiased high dimensional analysis to generate UMAPs from 10,000 downsampled cells from each stimulation and unstimulated condition. Cells colored as case and control are distributed uniformly throughout the plots, indicating no separation between the case and control samples **(Supplementary Figure 2C)**.

### HDStIM protocol and associated functions

#### Unsupervised K-means clustering followed by Fisher’s exact test

To identify the responding cells, for every SPC, expression values for all the response markers from single-cells from both the stimulated and unstimulated samples were subjected to K-means (K = 2) clustering. K-means clustering on the combined set of stimulated and unstimulated cells leads to a contingency table with the percentage of cells clustered in clusters 1 and 2 from both the experimental conditions. This contingency table was used to carry out a Fisher’s exact test, and SPCs that passed the Fisher’s exact test with a p-value < 0.05 were considered further. Thus, one with the higher stimulated/unstimulated ratio was labeled as the responding cluster of the two K-means clusters.

#### Diagnostic plots

HDStIM provides auxiliary functions for three types of diagnostic plots. For every SPC that passed Fisher’s exact test, users can plot 1) a stacked bar plot showing the contingency table used in Fisher’s exact test. This plot also mentions the p-value for Fisher’s exact test and the K cluster labeled as the responding cluster **(Supplementary Figure 3A)**. 2) an optional UMAP showing the separation of the cells into responding and non-responding classes. UMAPs are calculated on expression values of all the response markers from the single cells and colored as stimulated responding, stimulated non-responding, unstimulated responding, and unstimulated non-responding according to the cell selection method mentioned above **(Supplementary Figure 3B)**. 3) a distribution plot with stacked distributions for each response marker used in HDStIM. The stacked distribution plots show kernel density estimation for unstimulated, stimulated, and responding/mapped cells **(Figure 1C & 1D)**.

#### Ranking markers responsible for driving K-means clustering

HDStIM also provides a function to rank response markers responsible for driving the K-means clustering and subsequent partitioning of the cells into responding and non-responding. Under the hood, the function executes a random forest-based method, “Boruta” (Kursa, Miron B. and Rudnicki, Witold R., 2010). Boruta is an all-relevant feature selection algorithm that compares the importance scores obtained by features with the highest importance scored by the permuted (shadow) features by a two-sided test of equality. Features with significantly higher or lower importance than the permuted features are confirmed or rejected (p<0.05), with the remaining features deemed tentative. Utilizing the results from Boruta, the function generates plots with a mean marker importance score for each marker. The plots also show min and max marker importance scores as error bars and are colored to indicate if a particular marker was found significant in differentiating between the responding and non-responding cells against the shadow attributes. Besides Boruta, we also tried Regularized Random Forest and XGBoost Tree in R. However, the results were indifferent, or the compute time was longer than Boruta. Therefore, in favor of the robustness of an already established method and compute speed, we decided to adopt Boruta in our package. In this study, we have used importance scores calculated by Boruta and presented them either transformed or untransformed as heatmaps **(Figure 1 E, 1F, 2 B & 2 E)**.

#### Software availability, documentation, and reproduction of this study

HDStIM is available at CRAN at https://cran.r-project.org/package=HDStIM. The developmental version is available at https://github.com/niaid/HDStIM with an accompanying documentation website describing the included functions and their usage at https://niaid.github.io/HDStIM. To reproduce the figures presented in this article, data (including the FCS files from the pediatric dataset) and R scripts can be found at https://doi.org/10.5281/zenodo.6762108 and https://github.com/niaid/HDStIM-paper, respectively.

### Linear models to correlate fractions of responding cells with age

Outside the core functionality of the HDStIM package, we carried out linear modeling to correlate the fraction of responding cells with age in the pediatric dataset in R. We calculated fractions of responding cells per subject for cells deemed responding by HDStIM in unstimulated wells (baseline/tonic activation) and stimulated wells (after stimulation). We also took a difference (delta; in vitro stimulation) between after-stimulation and baseline fractions. For Figure 2 C, we regressed ranks of fractions of responding baseline and after stimulation cells on age, gender, and an interaction between the two. We used ranks of fractions than the fractions themselves to emulate non-parametric modeling. Of the three terms used as predictors, only age influenced the target.

For **Figure 2D** we carried out spline regression with knots on 8 and 14 years of age using linear polynomials. However, unlike linear regression for **Figure 2C**, we regressed responding baseline fractions only on age and only considered myeloid and T-cell populations for IFNa and TCR stimulation, respectively.

## Supplementary Information

**Supplementary Figure 1:**
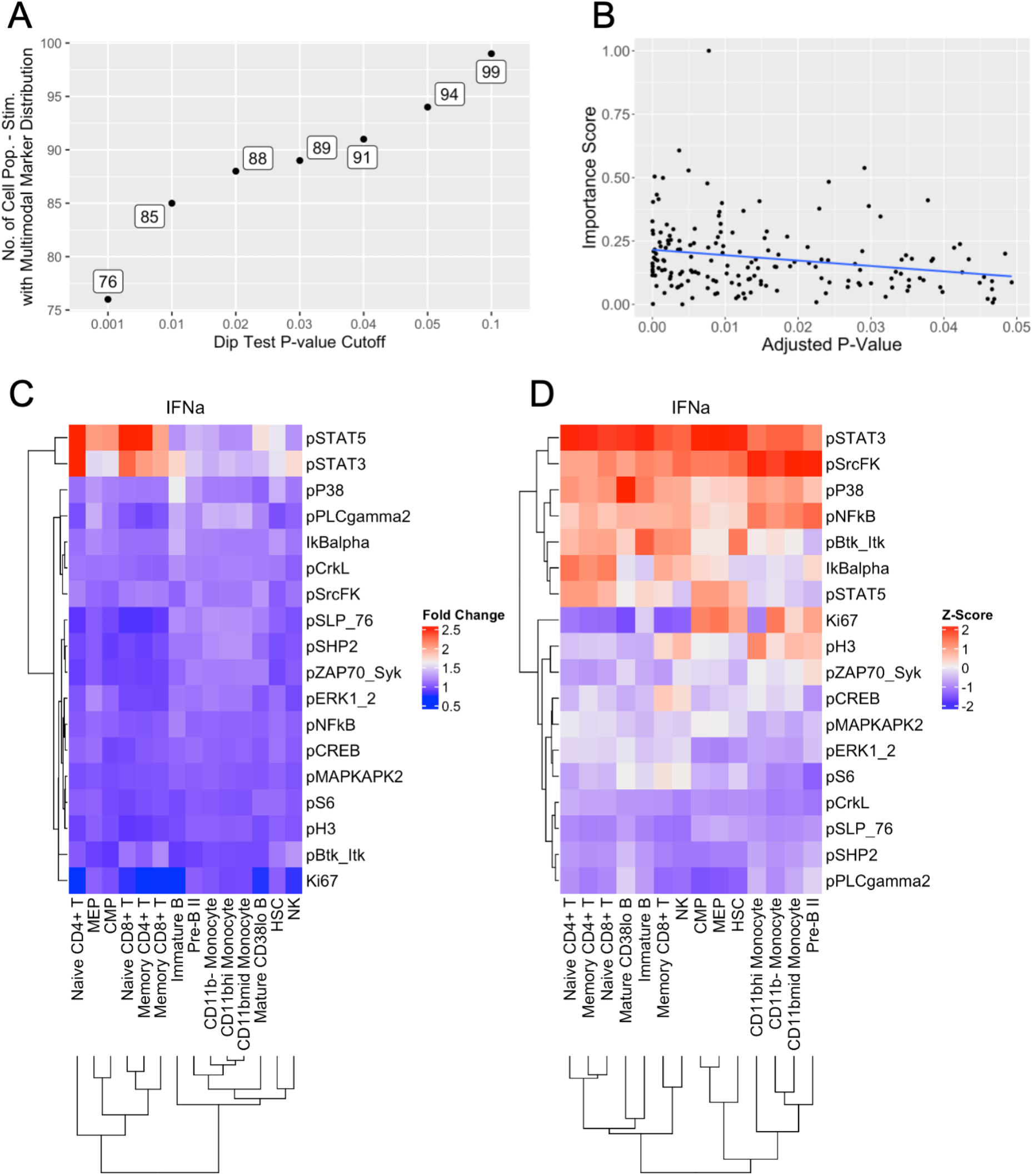
Estimation of multimodality in response markers and marker by marker analysis of pre-HDStIM data from the bone marrow dataset. A) Number of stimulation-cell population combinations (y-axis) with response markers with multimodal distributions at various cutoffs of dip test p-value (x-axis). B) Correlation between importance score calculated by HDStIM and adjusted p-values from individual marker analysis. Pearson correlation r = -0.22 with p-value = 0.003. Dots represent response markers from their respective stimulation-cell population combination. C) Fold change between unstimulated and stimulated samples for individual markers from IFNa stimulation. D) Z-scores calculated on mean expression values of individual response markers in the stimulated well from IFNa stimulation. Neither B nor C shows the pattern observed in Figure 1F. For comparison with Figure 1F in both B and C, only the responding cell populations identified by HDStIM are shown.

**Supplementary Figure 2:**
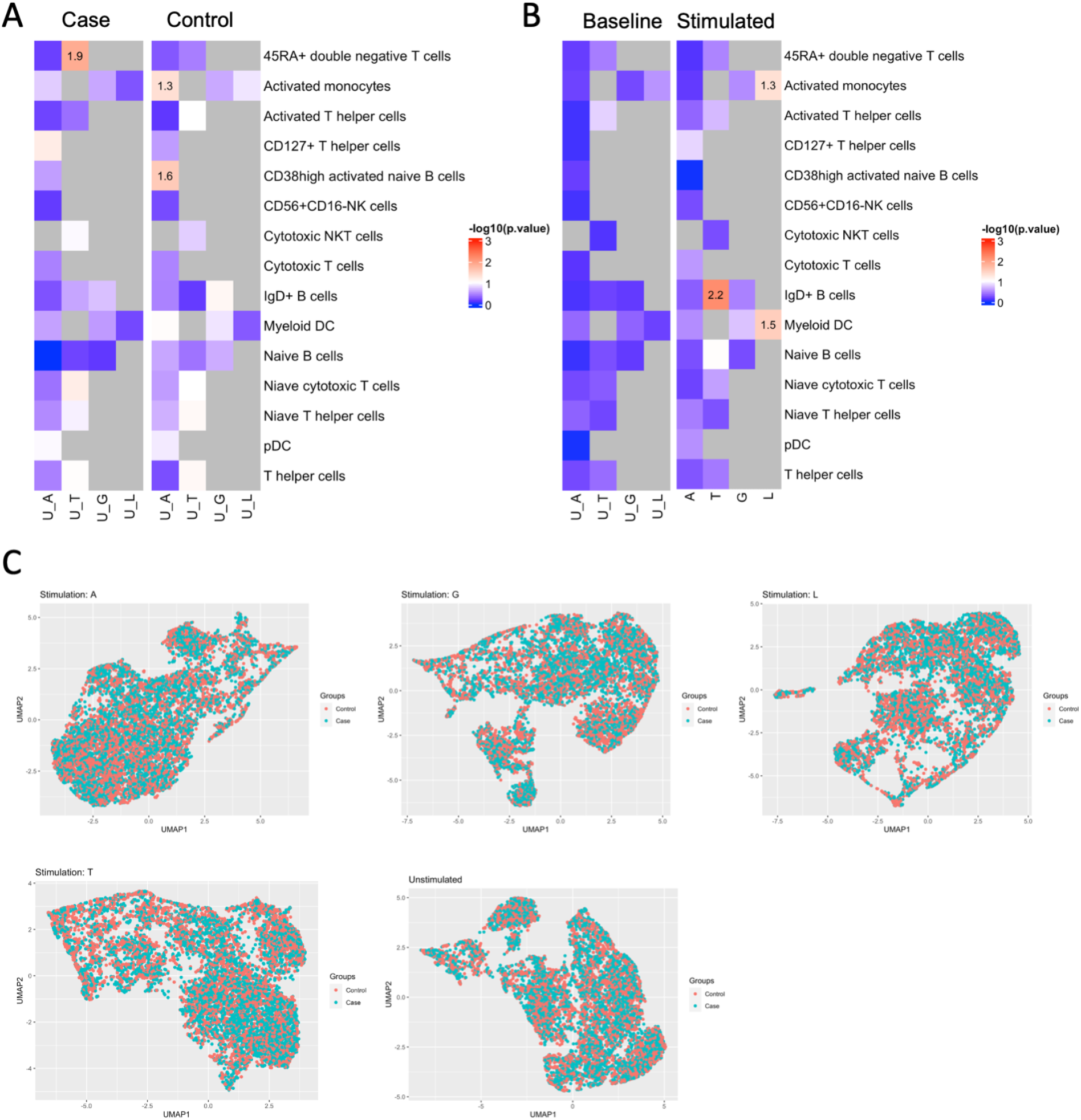
Effect of age on the fractions of responding cells at the baseline in disease (case) and healthy (control) samples from the pediatric cohort. A) Heatmap showing the -log10(p-value) of age from the linear regression where fractions of responding cells at the baseline were regressed on age, gender, and their interaction separately for disease (case) and healthy (control) samples. B) Heatmap showing the - log10(p-value) from the regression model of fractions of responding cells on case vs. control condition. C) A UMAP using the response markers on the 10,000 downsampled cells from each stimulated and unstimulated condition. Cells are colored red and blue for control and case samples, respectively.

**Supplementary Figure 3:**
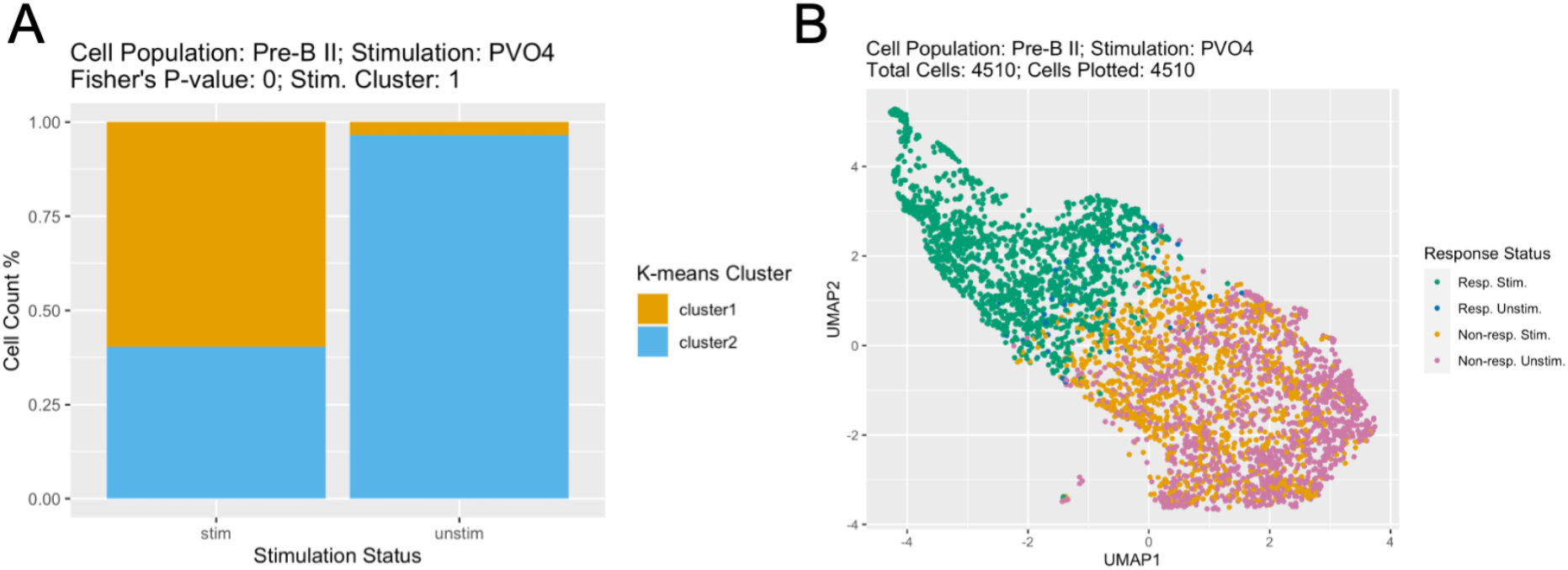
Diagnostic plots produced by HDStIM depicting the separation of responding from non-responding cells. A) A stacked bar plot showing the percentage of stimulated and unstimulated cells clustered in the two K-means clusters. This figure also represents the contingency table used to conduct Fisher’s exact test to select responding cells from the stimulated samples. B) A UMAP on all the cells from the Pre-B II cell population and PVO4 stimulation using the expression values of all the response markers. Cells colored in green and orange represent the responding and non-responding cells from the stimulated samples. Cells colored in blue and pink represent the responding and non-responding cells from the unstimulated samples.

**Supplementary Table 1:** Stimulation panel design for the bone marrow and pediatric datasets.

| Metal | Antigen | Marker Class* |
| --- | --- | --- |
| **Bone Marrow Dataset** |  |  |
| In115 | CD45 | type |
| La139 | CD45RA | type |
| Pr141 | pPLCgamma2 | state |
| Nd142 | CD19 | type |
| Nd144 | CD11b | type |
| Nd145 | CD4 | type |
| Nd146 | CD8 | type |
| Nd148 | CD34 | type |
| Nd150 | pSTAT5 | state |
| Sm147 | CD20 | type |
| Sm152 | Ki67 | state |
| Sm154 | pSHP2 | state |
| Eu151 | pERK1_2 | state |
| Eu153 | pMAPKAPK2 | state |
| Gd156 | pZAP70_Syk | state |
| Gd158 | CD33 | type |
| Gd160 | CD123 | type |
| Tb159 | pSTAT3 | state |
| Dy164 | pSLP_76 | state |
| Ho165 | pNFkB | state |
| Er166 | IkBalpha | state |
| Er167 | CD38 | type |
| Er168 | pH3 | state |
| Er170 | CD90 | type |
| Tm169 | pP38 | state |
| Yb171 | pBtk_Itk | state |
| Yb172 | pS6 | state |
| Yb174 | pSrcFK | state |
| Yb176 | pCREB | state |
| Lu175 | pCrkL | state |
| Cd110,111,112,114 | CD3 | type |
| **Pediatric Dataset** |  |  |
| Y89 | CD45 | type |
| Pr141 | CD7 | type |
| Nd142 | CD19 | type |
| Nd144 | pPLCg2 | state |
| Nd145 | CD4 | type |
| Nd146 | IgD | type |
| Sm147 | CD20 | type |
| Sm149 | CD25 | type |
| Nd150 | pSTAT5 | state |
| Eu151 | CD123 | type |
| Sm152 | AKT | state |
| Eu153 | pSTAT1 | state |
| Gd155 | CD27 | type |
| Gd156 | pP38 | state |
| Gd157 | CD24 | type |
| Gd158 | pSTAT3 | state |
| Tb159 | CD11c | type |
| Gd160 | CD14 | type |
| Dy163 | CD56 | type |
| Dy164 | IkBα | state |
| Ho165 | pCREB | state |
| Er166 | CD16 | type |
| Er167 | CD38 | type |
| Er168 | CD8 | type |
| Tm169 | CD45RA | type |
| Er170 | CD3 | type |
| Yb171 | pERK1/2 | state |
| Yb174 | HLA-DR | type |
| Lu175 | pS6 | state |
| Yb176 | CD127 | type |

\*Marker class: type = phenotyping marker; state = response marker.

**Supplementary Table 2:** Stimuli used in the bone marrow and pediatric datasets.

| No. | Symbol | Full Form |
| --- | --- | --- |
| Bone Marrow Dataset |  |  |
| 1 | BCR | B-cell Antigen Receptor |
| 2 | Flt3L | Flt-3 Ligand |
| 3 | GCSF | Granulocyte Colony Stimulating Factor |
| 4 | GMCSF | Granulocyte Macrophage Colony Stimulating Factor |
| 5 | IFNa | Interferon Alpha |
| 6 | IL3 | Interleukin 3 |
| 7 | IL7 | Interleukin 7 |
| 8 | LPS | Lipopolysaccharide |
| 9 | PMAiono | Phorbol Myristate Acetate |
| 10 | PVO4 | Pervanadate |
| 11 | SCF | Stem Cell Factor |
| 12 | TNFa | Tumor Necrosis Factor Alpha |
| 13 | TPO | Thrombopoietin |
| Pediatric Dataset |  |  |
| 1 | A | Interferon Alpha |
| 2 | G | Interferon Gamma |
| 3 | T | T-cell Receptor (TCR) |
| 4 | L | Lipopolysaccharide (LPS) |

**Supplementary Table 3:**
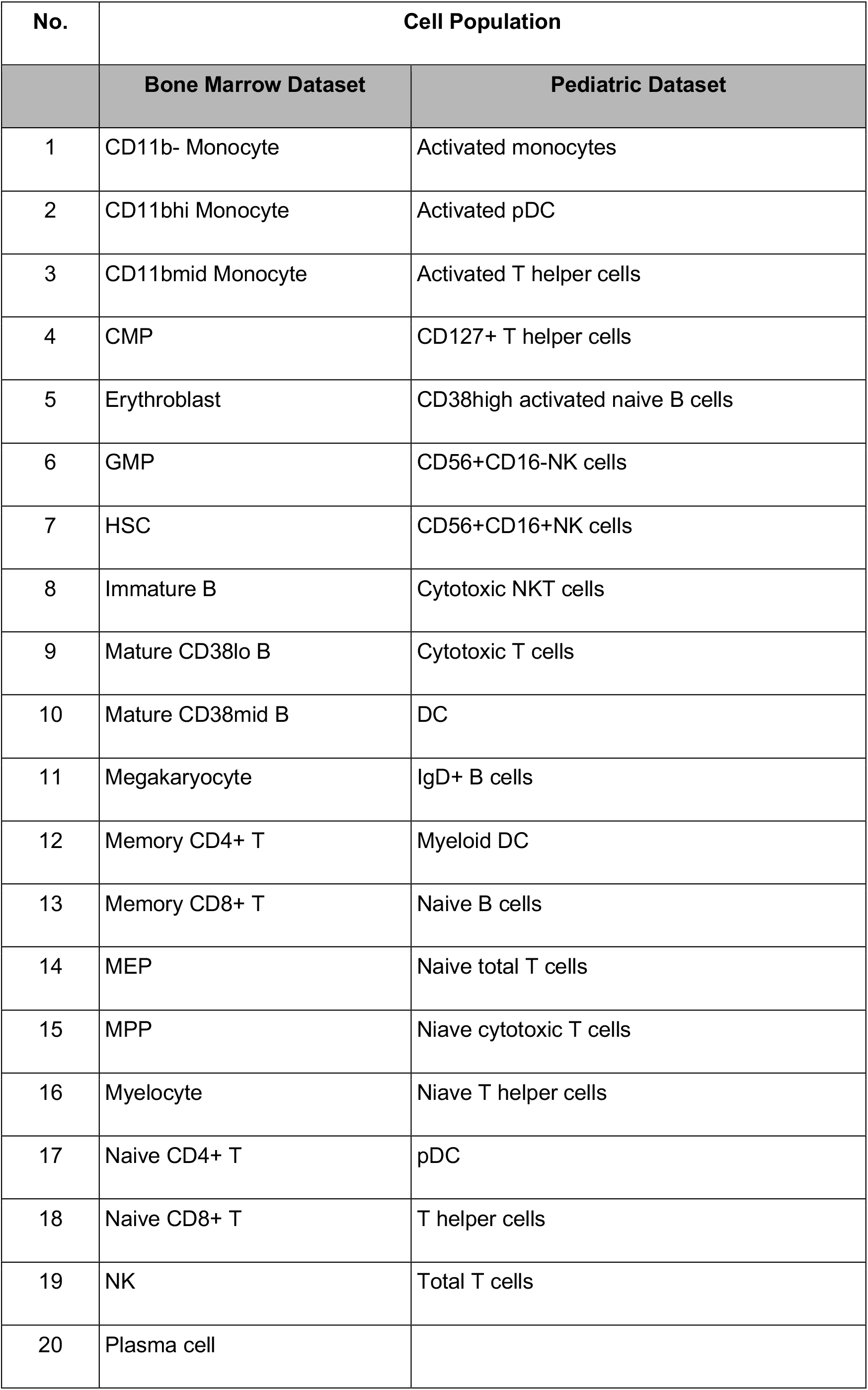
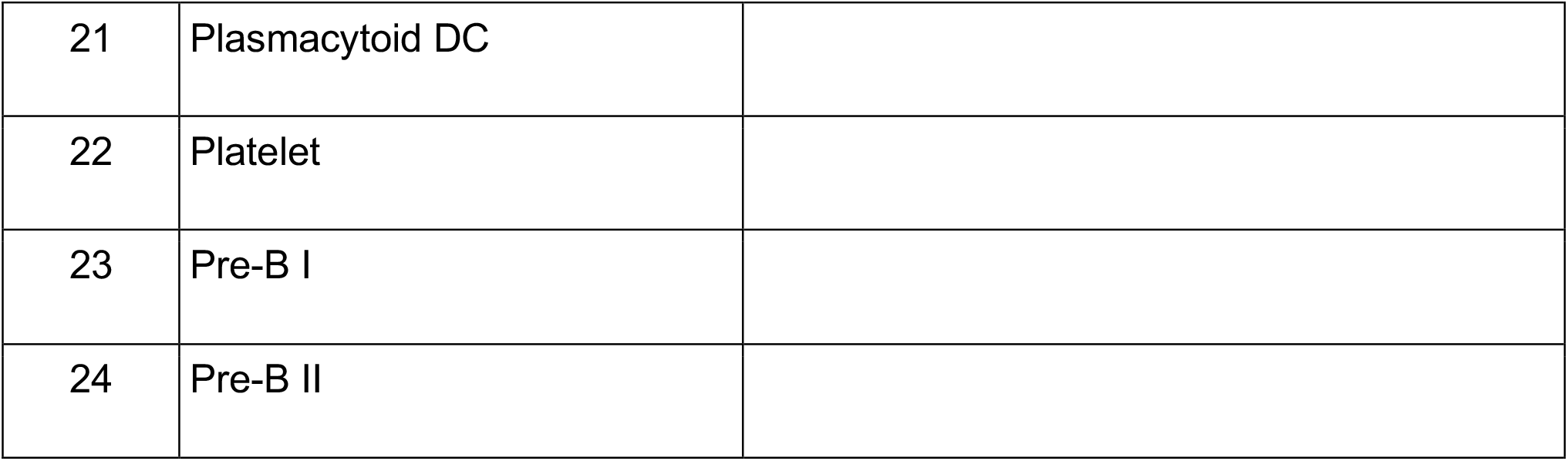
Major cell populations identified using phenotyping markers in the bone marrow and pediatric datasets.

## Ethics Statement

For this study volunteers were enrolled in 13-HG-0053, an NIH protocol approved and monitored by the National Human Genome Research Institute/NIH Institutional Review Board (Bethesda, Maryland, USA). All subjects provided informed consent prior to their participation in the study.

